# Identification of a novel anti-angiogenic regulatory sequence within the syndecan-3 extracellular core protein

**DOI:** 10.64898/2026.08.10.743887

**Authors:** Samantha Arokiasamy, Giulia De Rossi, Thomas C. Mosely, Sylvie Ricard-Blum, James R. Whiteford

## Abstract

Syndecans are transmembrane proteoglycans that regulate angiogenesis through both their glycosaminoglycan chains and core proteins. While roles for all four mammalian syndecans in new blood vessel formation are well established, it has more recently emerged that their extracellular core proteins contain discrete bioactive regulatory sequences capable of influencing cellular processes, including angiogenesis. We previously demonstrated that the syndecan-3 (SDC3) ectodomain possesses anti-angiogenic activity independent of its heparan sulphate chains. Here, we identified and characterised a novel anti-angiogenic sequence within the SDC3 ectodomain. Using recombinant truncation mutants, endothelial migration assays and peptide mapping, we localised activity to a discrete region of the extracellular domain and subsequently defined a conserved minimal nine amino acid peptide, QM111, that retained full biological activity. QM111 inhibited endothelial cell migration and angiogenic sprouting in both rat aortic ring and mouse choroidal explant models. Intrinsic disorder analysis revealed that QM111 resides within a region of comparatively reduced disorder, consistent with other syndecan regulatory sequences. This supports the concept that syndecan ectodomains contain conserved functional modules embedded within intrinsically disordered extracellular domains. QM111 did not induce inflammatory chemokine production, exhibited no detectable cytotoxicity, and retained substantial stability in human serum and vitreous humour. Finally, QM111 displayed anti-angiogenic activity comparable to the previously described syndecan-2-derived peptide QM107, with combination treatment producing more robust inhibition of angiogenesis. These findings identify QM111 as a novel endogenous anti-angiogenic peptide and support the concept that syndecan ectodomains are reservoirs of biologically active regulatory sequences with therapeutic potential. The work further establishes syndecan-derived peptides as a promising platform for the development of next-generation anti-angiogenic therapies.

## Introduction

Angiogenesis is essential for tissue growth, repair, and nutrient delivery. It occurs through the branching of new blood vessels from existing ones, a process governed by a balance of pro- and anti-angiogenic factors (Folkman, 2006). Key regulators, including vascular endothelial growth factor A (VEGFA) and fibroblast growth factor 2 (FGF2), drive endothelial cell proliferation, migration, and vessel formation. However, when dysregulated, angiogenesis contributes to disease. Excessive vascular growth fuels tumour progression in cancer, while fragile, leaky vessels in diabetic retinopathy impair vision (Cao et al., 2023). Understanding the molecular mechanisms underlying these processes, is crucial for developing targeted therapies to either enhance or inhibit angiogenesis as needed.

Syndecan-3 (SDC3), a member of the syndecan family of cell surface proteoglycans, has emerged as a key regulator of angiogenesis (Arokiasamy et al., 2020). While initially thought to be restricted to neural tissues (Kim et al., 1994), recent findings reveal its presence in endothelial cells, where it interacts with growth factors and extracellular matrix (ECM) components to influence vascular processes such as angiogenesis and permeability responses (De Rossi & Whiteford, 2013a; Jannaway et al., 2019). Structurally, SDC3 consists of a transmembrane core protein adorned with glycosaminoglycan (GAG) chains, namely heparan sulphate (HS) and chondroitin sulphate (CS), which facilitate interactions with ECM proteins, enzymes, and growth factors critical for angiogenesis (Asundi & Carey, 1995; Carey et al., 1992). Its extracellular domain modulates these interactions, while the intracellular domain links to signalling pathways that govern endothelial cell behaviour. Compared to the other family members, however relatively little is understood about the role of SDC3 in angiogenesis.

The interaction between pro-angiogenic growth factors such as VEGFA and FGF2 and heparan sulphate have been well characterised (Ashikari-Hada et al., 2005; Jemth et al., 2002; Robinson et al., 2006). It is therefore unsurprising that roles for all four syndecan family members have been identified in angiogenesis as co-receptors to promote growth factor signalling. Syndecan-1,-2 and −4 have all been implicated in VEGFA/VEGFR2 signalling through interactions with their HS chains (Chen et al., 2004; Corti et al., 2019; De Rossi et al., 2021; Rapraeger, 2013). However, it has also emerged that contained within syndecan extracellular core proteins are sequences that can modulate cell responses and processes, in particular angiogenesis (De Rossi & Whiteford, 2013b). The extracellular core proteins of all four family members exhibit anti-angiogenic properties independent of their heparan sulphate chains. In syndecan-1 a peptide sequence residing between L^93^ and E^120^ of the human sequence has been shown to inhibit angiogenesis in both *in vitro* and *in vivo* models (Beauvais et al., 2009). Similarly, a sequence residing between P^123^ and F^140^ of syndecan-2 has also been shown to be potently anti-angiogenic, acting through a novel VEGFA/VEGFR2 independent pathway involving the protein tyrosine receptor CD148 (Arokiasamy et al., 2025; Rossi et al., 2014; Whiteford et al., 2011). The syndecan-4 ectodomain also has anti-angiogenic properties although the precise peptide sequence has yet to be mapped, however, it would likely be associated with a conserved NXIP motif known to be important for driving cell adhesion responses (De Rossi et al., 2021; Whiteford & Couchman, 2006). Previously we have shown that the extracellular core protein of SDC3 also inhibits angiogenesis although the sequence and indeed mechanism are not known (De Rossi & Whiteford, 2013a).

In this study, we investigated the molecular basis of the anti-angiogenic activity associated with the SDC3 extracellular domain. Using recombinant proteins, peptide mapping and functional angiogenesis assays, we identified a novel bioactive sequence within the SDC3 ectodomain that suppresses endothelial cell responses and angiogenesis. Our findings establish SDC3 as a further source of extracellular matrix-derived regulatory peptides with antiangiogenic activity and potential therapeutic application.

## MATERIALS AND METHODS

### Cell culture

Human umbilical vein endothelial cells (HUVECs) were obtained from Promocell (Cat# C-12203) and were grown in Endothelial Cell Growth Medium 2 (ECGM2, Promocell Cat# C-22111) on tissue culture flasks and dishes coated with 0.1% gelatin from porcine skin (Merck Life Science Cat# G1890), at 37°C, 10% CO_2_. Human retinal pigment epithelial cells (H1RPE7 cell line) were obtained from HPA culture laboratories (Public Health Laboratories, UK Cat# 09061602, RRID:CVCL_2429) and were grown in Ham’s F12 medium (ThermoFisher Scientific, Cat#31765035) supplemented with 20% heat-inactivated Fetal Bovine Serum (FBS), 2 mM Glutamine (ThermoFisher Scientific, Cat# 25030081) and a selection antibiotic 1 μg/ml puromycin (ThermoFisher Scientific, Cat# A1113803). Endothelial cells isolated from brain tissue derived from a mouse with endothelioma (bEND.3 cells) were grown in Dulbecco’s Modified Eagle Medium (DMEM, ThermoFisher, Cat#21885-025) supplemented with 10% FBS.

### Expression and purification of full-length and truncated forms of murine SDC3 ectodomain

The entire length (A^45^-L^380^) of the murine SDC3 ectodomain (S3ED) was cloned into the pET41 expression vector (Merck, Cat# 70566) as described in (De Rossi & Whiteford, 2013a). The recombinant proteins were expressed as glutathione-S-transferase (GST) fusion proteins with GST located at the N-terminus. To generate the truncated forms of S3ED; namely S3EDA^45^-A^194^, S3EDP^195^-L^380^, S3EDE^92^-V^310^, S3EDE^151^-A^221^ and S3EDP^195^-A^221^, the corresponding coding sequences with additional 3’ stop codons were amplified by PCR using the primers described in (Supplemental Table 1). PCR products were then digested with *EcoRI* and *HIndIII* and ligated into the equivalent sites of pET41 using standard procedures. S3ED and its truncated forms were expressed in and purified from cultures of *E. coli* strain BL21(DE3) (Merck Cat#69450) grown to an OD600 of 0.4 prior to the addition of 0.1 M Isopropyl β-D-1-thiogalactopyranoside (IPTG) and subsequent outgrowth for 4 hours. Purifications were performed using glutathione Sepharose 4B (Merck Cat#GE17-0756-01) according to the manufacturer’s instructions.

### Syndecan derived peptides

The peptides QM111M (PPATATVADVRTTGIQGMLPLPLTTAA), QM111H (PPFTATTAVIRTTGVRRLLPLPLTTVA), QM111 (LPLPLTTVA), QM111S1 (ALLLPTVTP), QM111S2(PPALLVTLT), QM107 (PAEENTNVYTEKHSDSLF) were all synthesized by Cambridge Peptides UK. Scaled synthesis (∼2 g) of the QM111 peptide was performed by Bachem Peptides. All peptide preparations were obtained at >95% purity as determined by HPLC.

### Prediction of intrinsic disorder in peptides

Protein disorder was predicted in the syndecan peptides using IUPred2A web interface (Short disorder) that allows to identify disordered regions (https://iupred2a.elte.hu/) (Mészáros et al., 2018). ANCHOR predicts binding regions that are disordered but can undergo disorder-to-order transition upon binding to partners (Dosztányi, 2018).

### *Ex vivo* angiogenic sprouting assays

Thoracic aortas were isolated from male Wistar rats (∼200 g, Charles River UK). Surrounding adipose tissue and side branches were removed prior to the sectioning of aortas into rings of <1 mm thickness. Choroidal explants were prepared from male C57BL/6J mice (3–4 weeks old, Charles River UK). Eyes were punctured and the cornea, iris, lens and retina removed before the remaining choroid was dissected into approximately 1 mm³ explants. Following dissection, aortic rings and choroidal explants were incubated overnight at 37°C in Opti-MEM (ThermoFisher Scientific, Cat#31985070) before being embedded in 0.15 ml of 1 mg/ml collagen I (Millipore, Cat#06-115) prepared in E4 medium (ThermoFisher Scientific, Cat#12800-017) in 48-well plates (Corning). Collagen gels were allowed to polymerise for 30 min at 37°C before being overlaid with 200μl Opti-MEM supplemented with 1% heat-inactivated FBS. Angiogenesis was stimulated with VEGFA (VEGF_165_, R&D Systems, Cat#293-VE) at 10 ng/ml for rat aortic rings and choroidal explants. Experimental treatments were added at the indicated concentrations, and culture medium was refreshed every three days. Microvascular sprouts were counted using an Olympus IX81 inverted microscope after seven days and expressed as the number of sprouts per aortic ring or choroidal explant.

### Scratch wound migration assay

Confluent monolayers of ECs were scratched with a pipette tip to generate a ‘wound’. Cells were then washed once with PBS prior to the addition of growth medium supplemented with the treatments as described. Scratch wounds were monitored by time-lapse microscopy using an Auto LCI incubator imaging system (World Precision Instruments) and wound closure calculated using the Auto LCI imaging software (World Precision Instruments).

### Chemokine proteome profiler assay and CXCL8 and CCL2 measurements by ELISA

The Proteome Profiler Human Chemokine Array Kit (Biotechne, Cat# ARY017) was used to determine the levels of 31 chemokines in HUVEC conditioned media harvested 4h after the treatments indicated. Arrays were used as per the manufacturer’s instructions and imaged on an Azure C600 bioimager. Densitometry of chemokine spots was performed using Image J. Quantification of CXCL8 and CCL2 was performed using Quantikine® ELISA Kits as per the manufacturer’s instructions (CXCL8 Cat# HS800, CCL2 Cat# DCP00 both from R&D Systems).

### *In vivo* peritonitis model

All animal care and experimental protocols were conducted at the William Harvey Research Institute, Queen Mary University of London, UK under the UK legislation for animal experimentation (UK Home Office license number PPL: P873F4263) and in agreement with the UK Home Office Animals Scientific Procedures Act 1986. Animal studies are reported in compliance with the ARRIVE guidelines (Lilley et al., 2020; Percie du Sert et al., 2020).

Peritonitis was induced in male C57BL/6 mice between 8 and12 weeks old by injecting 1 µg of either LPS (lipopolysaccharides from *E. coli*, Sigma-Aldrich, Cat #297-473-0) or/and 50 µM QM111 intraperitoneally (i.p.) in 0.5 ml sterile PBS. After 4 hours, mice were sacrificed by cervical dislocation and peritoneal cells collected by lavage of the peritoneum with 6 ml of PBS supplemented with 0.25% w/v BSA. Harvested fluid was centrifuged at 250 x g for 5 min and cell pellets were resuspended in FACS-buffer (PBS supplemented with 1% v/v heat-inactivated FBS) ready for fluorescent labelling.

### Immunofluorescence labelling of leukocytes and flow cytometry

Leukocytes were identified by their size (FSC) and granularity (SSC), CD45.2 positive staining and DAPI negative staining. Neutrophils and macrophages were identified based on Ly6G- and F4/80-high staining respectively. Cells were fluorescently labelled with various fluorochromes using monoclonal antibodies recognising CD45.2 (BioLegend, Cat. #109828), Ly6G (BioLegend, Cat. #127609), and F4/80 (0.05 μg/ml, BioLegend, Cat. #123114) and using DAPI to assess cell viability (Sigma-Aldrich, Cat. #D9542) for at least 1 h at 4 °C along with 0.5 μg FcγIII/II receptor blocking antibody (Fc-block™, BD Biosciences, Cat#553142) to prevent unspecific Fc-receptor binding. Antibody-labelled cells were washed twice with FACS buffer before being resuspended in 200 µl cold FACS buffer to achieve a single cell suspension. Following this, all samples were kept on ice. DAPI (1 μg/ml), a fluorescent dead cell nuclear marker, was added prior to analysis by flow cytometry. Samples were run on a BD LSR-Fortessa flow cytometer (BD Biosciences) using FACSDIVA software. Multi-colour fluorescence overlap was compensated for by using single-stained BDTM CompBeads (BD Biosciences Cat#552844). At least 20,000 events were acquired per sample. Data were analysed with FlowJo analysis software (Treestar).

### Generation of a polyclonal antibody to QM111

A polyclonal antibody against QM111 was generated commercially by Covalab (Bron, France) using two New Zealand White rabbits. QM111 was conjugated to Keyhole Limpet Hemocyanin (KLH) and administered with complete Freund’s adjuvant on days 0, 21, 42 and 63. Preimmune serum was collected prior to the initial immunisation (day 0), with test bleeds obtained on days 53 and 74. A terminal bleed was performed on day 88, and antisera were used for subsequent ELISA analyses.

### Measurement of QM111 levels in biological samples by ELISA

QM111 concentrations were quantified by ELISA using standard procedures. Briefly, QM111 calibrators and experimental samples were coated onto 96-well MaxiSorp microtitre plates (ThermoFisher Scientific, Cat#442404) by incubation overnight (16 h) at 4°C. Plates were washed three times with PBS before blocking with 100 μl of 1% (w/v) BSA in PBS for 30 min at room temperature. Following a further series of PBS washes, primary antibody (crude antiserum) diluted in PBS (100 μl) was added to each well and incubated for 1 h at room temperature. Wells were then washed and incubated with horseradish peroxidase-conjugated secondary antibody (100 μl, 1:500 dilution, ThermoFisher Cat#31460) for a further 1 h at room temperature. After a final wash, bound antibody was detected using TMB One Solution (Promega, Cat#G7431) according to the manufacturer’s instructions. The reaction was terminated after 5 min by the addition of 0.1 M HCl, and absorbance was measured at 450 nm using a Spectra MR plate reader (Dynex Technologies).

### Testing QM111 peptide stability in human serum and vitreous humour

Vitreous humour was extracted from enucleated pig’s eyes and diluted in an equal volume of PBS. Peptide samples (25 µl) diluted in PBS were added to 25 µl of vitreous humour or human serum and incubated at 37°C for the times indicated. Reactions were stopped by the addition of 100 µl of 4% w/v paraformaldehyde in PBS. Samples were then applied to 96 well plates and the peptide detected by ELISA as described above.

### Human serum samples

Human serum samples were collected from healthy donors (21-50 years of age, male and female.) All healthy volunteers gave written, informed consent to blood collection and the procedure was approved by the Queen Mary Ethics of Research Committee (QMERC2014.61).

### Lactate dehydrogenase cytotoxicity assay

Cytotoxicity was measured using the ThermoFisher™ Pierce™ LDH Cytotoxicity Assay Kit (Cat# 13454269) as per the manufacturer’s instructions.

### Statistical analysis

Statistical analyses were performed using GraphPad Prism (version 11.0.2; GraphPad Software, San Diego, CA, USA). Data are presented as mean ± SEM unless otherwise stated. Comparisons between multiple groups were performed using one-way ANOVA followed by Dunnett’s multiple comparisons test unless otherwise indicated in the figure legends. Differences were considered statistically significant at P < 0.05.

## RESULTS

We generated a fusion protein comprising the full length of the murine SDC3 extracellular core protein sequence (A^45^-L^380^) N-terminally fused to GST (**Figure 1A**) and expressed and purified this protein from bacterial lysates (De Rossi & Whiteford, 2013a). Confirming previous observations, it strongly inhibited (>60%) angiogenic sprout formation from rat aortic ring explants which was not the case in rings treated with GST alone (**Figure 1B**). We next set out to identify the regions of the murine SDC3 sequence that were responsible for its anti-angiogenic properties. We first generated two truncated ectodomains of SDC3, one comprising the N-terminal sequence A^45^-A^194^ and the other the C-terminal sequence P^195^-L^380^ (**Figure 1A**) and examined their effects on sprout formation from rat aortic ring explants. Whilst the N-terminal protein S3EDA^45^-A^194^ did not exhibit any inhibitory effects on angiogenesis in this model, the C-terminal portion exhibited a partially inhibitory effect (∼30%) as compared to the GST control, suggesting that the regulatory sequence might reside within this sequence (**Figure 1B**). To map this further we adopted a reductive approach whereby the S3ED sequence was truncated to progressively smaller protein sequences (summarized in **Figure 1A**). These were tested in endothelial cell (HUVEC) scratch wound assays. As expected, full length S3ED inhibited EC migration (∼50%) as compared to vehicle and GST controls. S3EDA^45^-A^194^, corresponding to the N-terminal portion of S3ED, did not inhibit EC migration consistent with it also not inhibiting angiogenic sprout formation. In contrast, inhibition (∼25%) was observed in HUVECs treated with S3EDP^195^-L^380^, and these properties were retained in a truncated protein encompassing sequences of both the N and C-terminus of S3ED (S3EDE^92^-V^310^). Further truncations revealed that a 27 amino acid sequence lying between P^195^ and A^221^ of murine S3ED inhibit EC migration (∼75%) (**Figure 1 C and 1D**). To confirm the importance of this region for the anti-angiogenic activity of S3ED, we generated a deletion mutant lacking residues P^195^-A^221^ (S3EDΔP^195^-A^221^; **Figure 1E**). This deletion abolished the inhibitory effect of S3ED on endothelial cell migration, restoring migration to levels comparable with PBS and GST controls, whereas full-length S3ED inhibited migration by ∼25% (Figure 1F). These data identify residues P^195^-A^221^ as essential for the anti-angiogenic activity of the SDC3 ectodomain.

**Figure 1:**
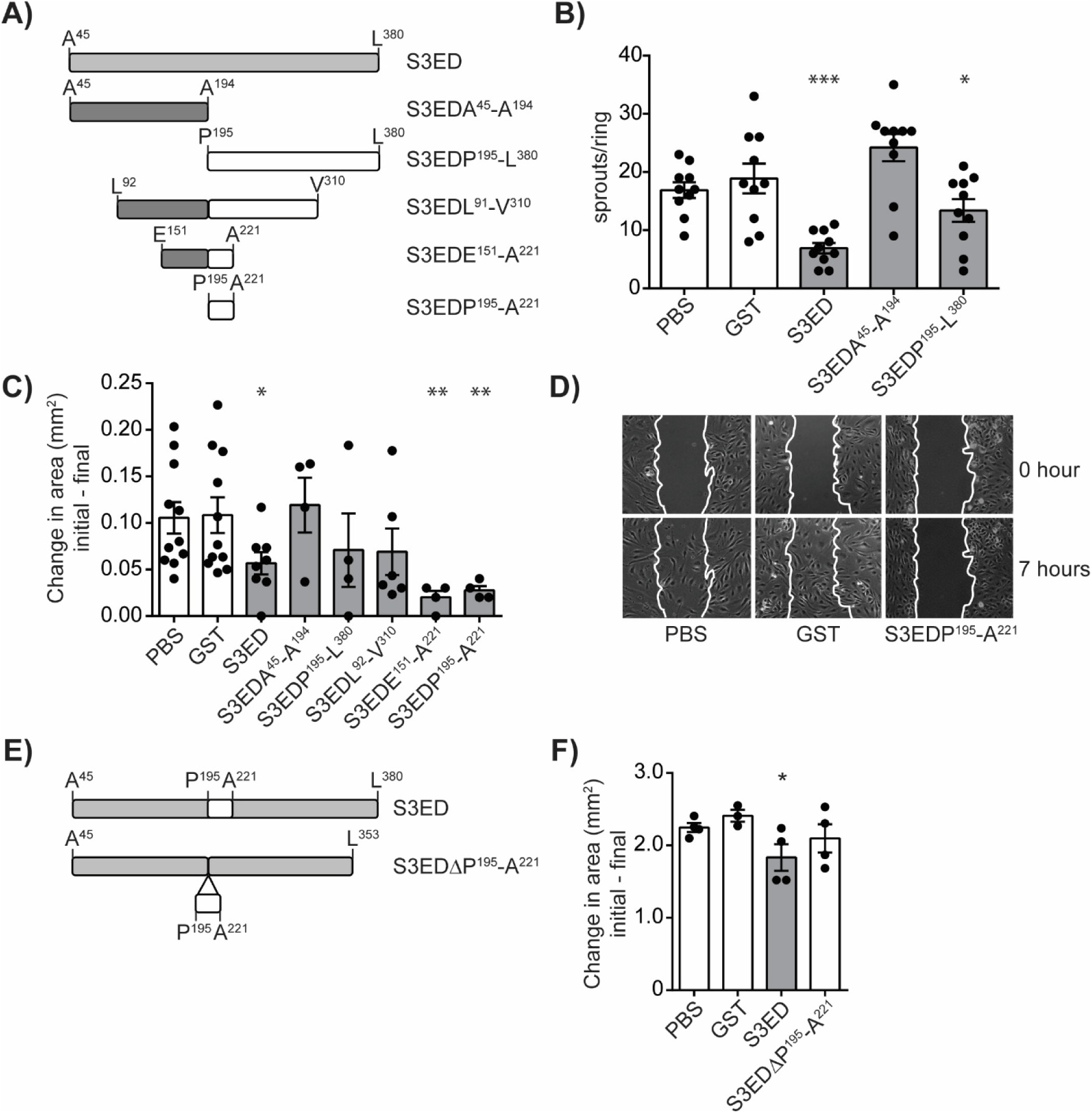
The anti-angiogenic properties of the murine SDC3 extracellular core protein require the sequence between P^195^ and A^221^. **A)** Schematic showing the strategy for identifying which region of the SDC3 extracellular core protein is required for the inhibition of angiogenesis. Coding sequence from murine SDC3 was translationally fused at the N-terminus to GST. **B)** The anti-angiogenic properties of S3ED reside in the C-terminal half of the SDC3 ectodomain between P^195^ and L^380^. Rat aortic rings were treated with 0.5 µM of the proteins indicated and the number of sprouts counted 7 days after seeding (n=∼10 rings/condition from aortas isolated from 3 animals, *p≤0.05, ***p≤0.001). **C)** HUVEC scratch wound migration assays reveal that the anti-angiogenic properties of S3ED map to a 27 amino acid sequence between P^195^ and A^221^ in murine SDC3. Scratches were performed on confluent HUVEC monolayers after which 0.5 µM of the proteins (0.5 µM) indicated were added. Images were captured after 0 and 7 hours and the change in area calculated (n=4-12, **p≤0.01). **D)** Micrographs of HUVEC monolayers comparing control (GST) treatment with the S3EDP^195^-A^221^ fusion protein. Wound closure is inhibited by S3EDE^195^-A^221^. **E)** Schematic showing the deletion of the P^195^-A^221^ anti-angiogenic region of the SDC3 ectodomain to generate S3EDΔP^195^-A^221^. **F)** S3EDΔP^195^-A^221^ does not inhibit HUVEC migration when compared to S3ED indicating the requirement for P^195^-A^221^ for the anti-angiogenic properties of the SDC3 ectodomain (n=4, *p≤0.05).

Inhibition of angiogenesis is a strategy for treating several diseases including cancer and neovascular eye diseases. We had previously shown that a peptide derived from the SDC2 ectodomain has the potential to be utilized as an anti-angiogenic therapy for neovascular eye diseases (Arokiasamy et al., 2025). Peptide therapies offer significant advantages over larger biologics due to simpler, less expensive manufacturing, superior tissue penetration, lower immunogenicity, and enhanced structural stability. For this reason, we next sought to see if peptides corresponding to P^195^-A^221^ of murine SDC3 (QM111M) and the corresponding region of human SDC3 (QM111H) could inhibit neovascularisation. The human sequence bears 66.7% identity and 81.5% similarity to the murine sequence (**Figure 2A**). Both peptides inhibited HUVEC cell migration to a similar degree (∼50%) and angiogenic sprout formation from rat aortic rings (>70% inhibition) indicating that this biological activity is conserved between species and retained in peptide form (**Figure 2B and 2C**). Comparison of the murine and human sequences showed that the peptides were highly conserved, particularly in the 9 C-terminal residues which were identical except for one amino acid residue (A in the murine sequence and V in the human sequence, **Figure 2D**). We therefore generated a peptide comprised of the human version of these 9 amino acids (QM111) and two scrambled controls (QM111S1 and QM111S2). QM111 retained the inhibitory properties of the larger QM111H in HUVEC scratch wound assays where it inhibited cell migration by 50% (**Figure 2E**) whereas the scrambled forms did not (**Figure 2F**).

**Figure 2:**
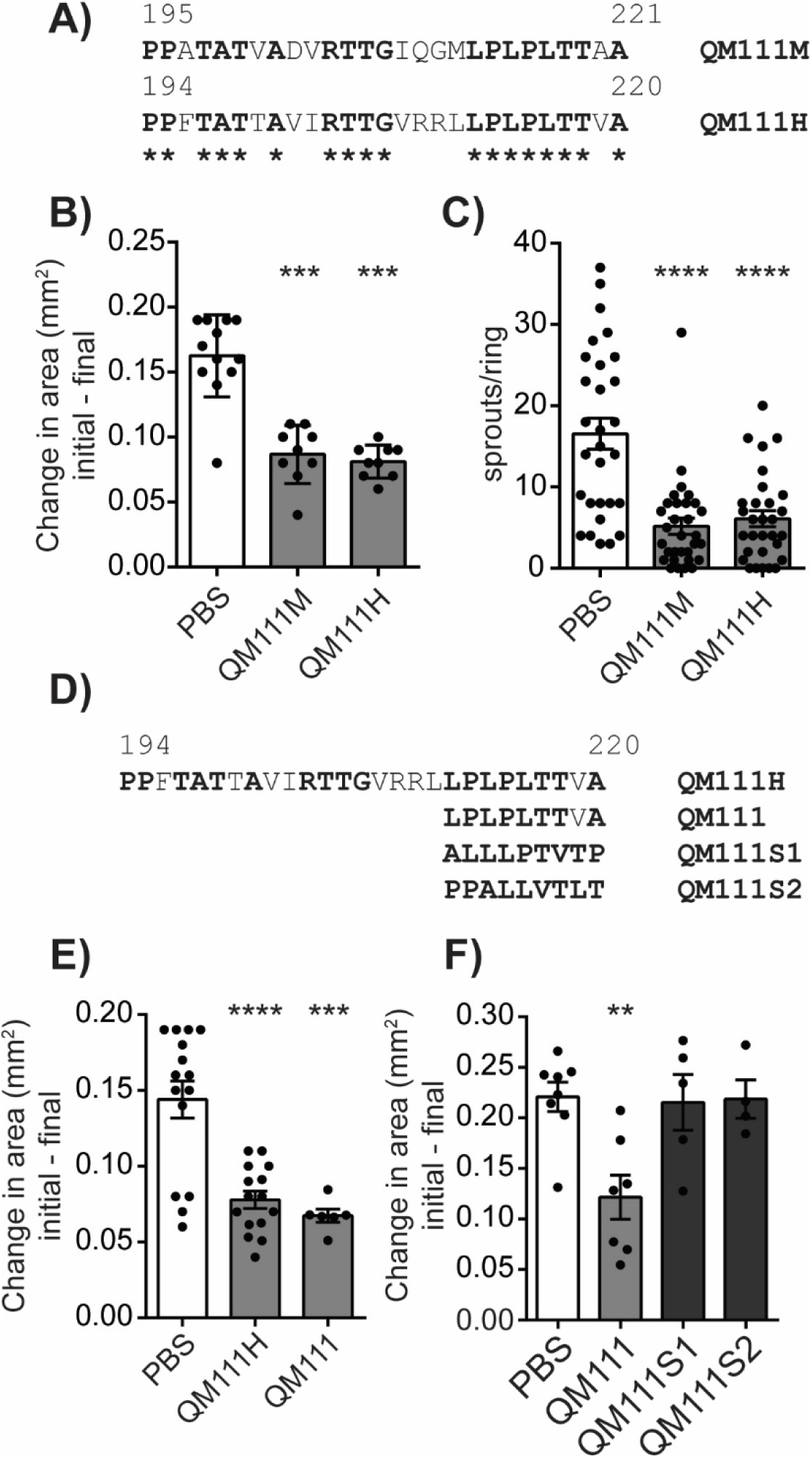
Identification of the minimal anti-angiogenic peptide sequence from SDC3. **A)** Comparison of the mouse and human peptide sequences corresponding to the P^195^-A^221^ region of murine SDC3. **B)** Peptides corresponding to the murine SDC3 sequence (P^195^-A^221^, QM111M) and the equivalent human sequence (P^194^-A^220^) were synthesized and tested for their anti-angiogenic potential in a scratch wound assay on confluent HUVECs. Cells were treated with 0.5 µM of each peptide and both peptides exhibited similar levels of inhibition of cell migration as compared to the PBS control (N=∼9, ***p≤0.001). **C)** Both QM111M and QM111H also showed equivalent inhibition of angiogenesis when applied to the rat aortic rings model (n=∼15, ****p≤0.0001). **D)** A peptide of 9 amino acid residues corresponding to the conserved C-terminus of murine and human sequences was synthesized based on the human sequence, QM111, together with two scrambled versions (QM1111S1 and QM111S2). **E)** QM111 showed equivalent inhibition of HUVEC cell migration to the parent QM111H peptide in scratch wound assays (n=∼15, ***p≤0.001, ****p≤0.0001. **F)** The scrambled forms of QM111 do not inhibit cell migration (n=7, **p≤0.01).

Syndecan ectodomains been experimentally shown to be intrinsically disordered. The ectodomain of human SDC3 contains about 69% of random coil and is the most disordered together with the ectodomain of syndecan-1 (Gondelaud et al., 2021; Ricard-Blum & Couchman, 2023;). We therefore used the IUPred2 algorithm to analyse intrinsic disorder in the SDC3 ectodomain and found that the QM111 sequence maps to a region with lower predicted intrinsic disorder than the surrounding ectodomain (**Figure 3 A and 3B**). Interestingly, the regulatory sequences identified within syndecan-1 (Beauvais et al., 2009) and syndecan-2 (Arokiasamy et al., 2025) also reside within regions of comparatively reduced predicted intrinsic disorder (**Supplemental Figure 1A, Figure 3C**). Although predictive, this observation suggests that bioactive sequences within syndecan ectodomains might be more structured. However, these sequences are embedded in highly flexible ectodomains that adopt several conformational ensembles, and seem to be partially masked, and even buried, in some conformations *in vivo* (Gondelaud et al., 2021). Thus, the QM111 sequence might not be available *in vivo* in all the conformations of SDC3.

**Figure 3:**
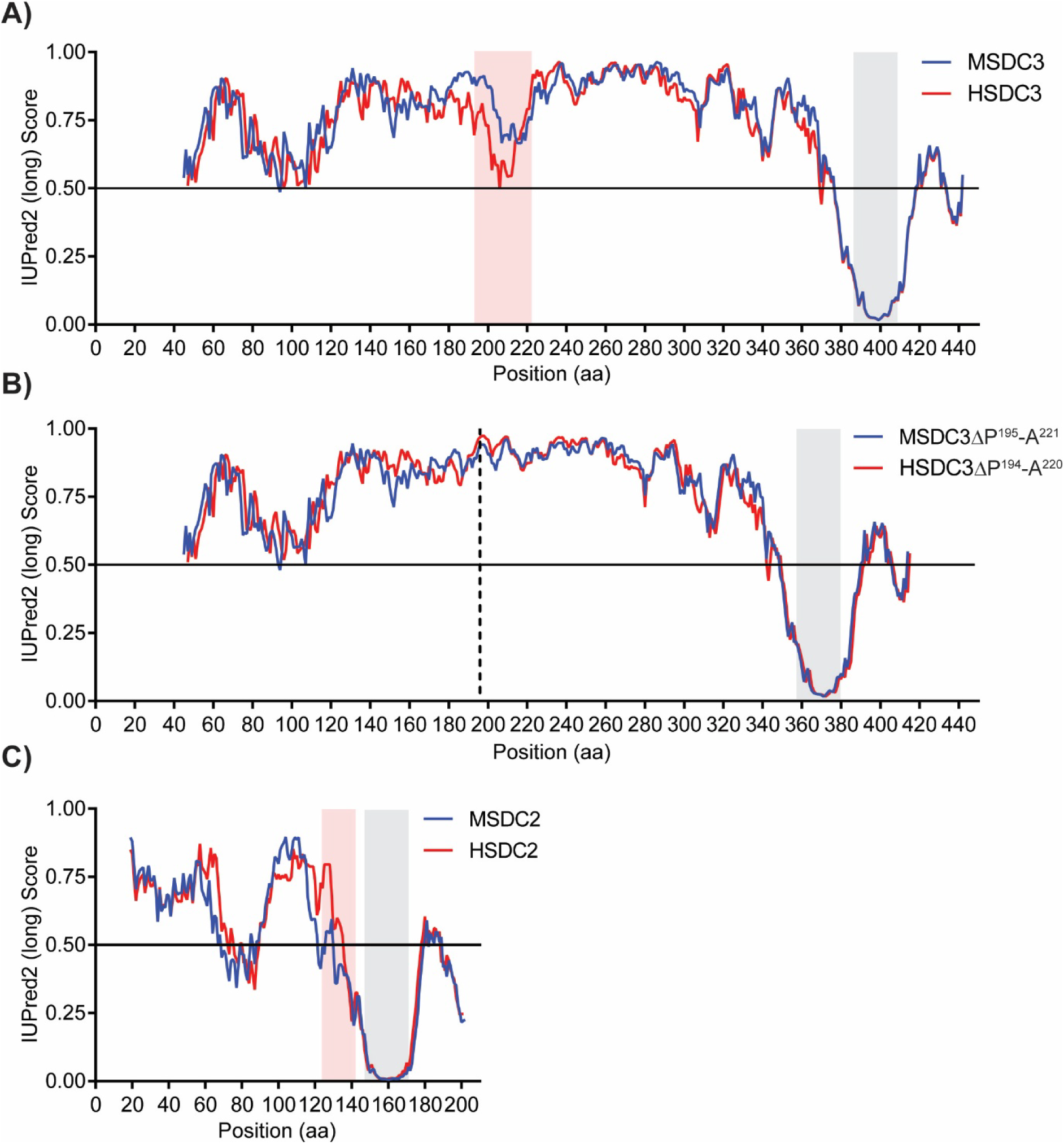
The anti-angiogenic sequence of SDC3 is predicted to reside in a less intrinsically disordered region of its ectodomain. **A)** Predicted intrinsic disorder profile for the entire length of the human (red) and mouse (blue) SDC3 amino acid sequence. The transmembrane region is highlighted by a grey box and the anti-angiogenic sequence in a red box. **B)** Predicted intrinsic disorder profile for the mutant form of mouse and human SDC3 lacking the anti-angiogenic region (MSDC3ΔP^195^-A^221^, HSDC3ΔP^194^-A^220^). **C)** Predicted intrinsic disorder profile for the full-length human and mouse SDC2 amino acid sequence indicating that the anti-angiogenic region is also predicted to fall in a less disordered region of the ectodomain.

The QM111 peptide, that recapitulates the anti-angiogenic activity of SDC3, has potential therapeutic applications, and was also analysed as an isolated sequence because disorder is dependent on the molecular context. QM111 was predicted to be entirely disordered (100%), whereas the murine and human QM111M and QM111H peptides were predicted to be intrinsically disordered at the N- and C-termini only, with a global disorder of 67% and 44% respectively (**Supplemental figure 2A and B**). The QM111, Q111M and Q111H peptides might not have the same flexibility and might adopt different conformations, which could modulate their interaction repertoire and/or their affinity for their partners. This could explain the slight differences observed between the peptides in the biological assays. The prediction of disordered protein binding regions in these peptides showed that the disordered sequences that might undergo a disorder-to-order transition upon binding to a protein partner. No protein binding disordered region was predicted in QM111 and in QM111H. However, a putative binding region was found in the C-terminus of the QM111M (**Supplemental figure 3**).

Having shown the anti-angiogenic potential of QM111 we sought to assess whether it could potentially be a viable therapeutic. A key consideration is whether it could cause off target adverse inflammatory responses. To test this, we first investigated whether HUVECs treated with QM111 synthesized higher levels of chemokines. Using a human proteome profiler chemokine array, we analysed conditioned media from cells treated with an inflammatory cocktail (TNF, IL1β and LPS), PBS, QM111 and the inflammatory cocktail with QM111. As expected, the inflammatory cocktail stimulated chemokine production, notably CXCL1, CXCL8, CCL2 and CXCL7, PBS did not elicit a response and importantly neither did QM111 (**Figure 4A and B**). The inflammatory response in cells treated with both the inflammatory cocktail and QM111 was equivalent to that of cells treated with the inflammatory cocktail alone. To quantify this further we measured CXCL8 and CCL2 levels by ELISA in conditioned media after the above treatments and observed the same result. Cells treated with the inflammatory cocktail showed substantially elevated chemokine levels (8-fold increase for CXCL8, 3-fold increase for CCL2) which was not the case with cells treated with either PBS or QM111 (**Figure 4C**). Finally, we used an *in vivo* peritonitis model and observed strong neutrophil ingress into the peritoneal cavity and reduced macrophage numbers in peritoneal lavage after LPS injection but not after QM111 injection where levels remained equivalent to the PBS control (**Figure 4D and 4E**).

**Figure 4:**
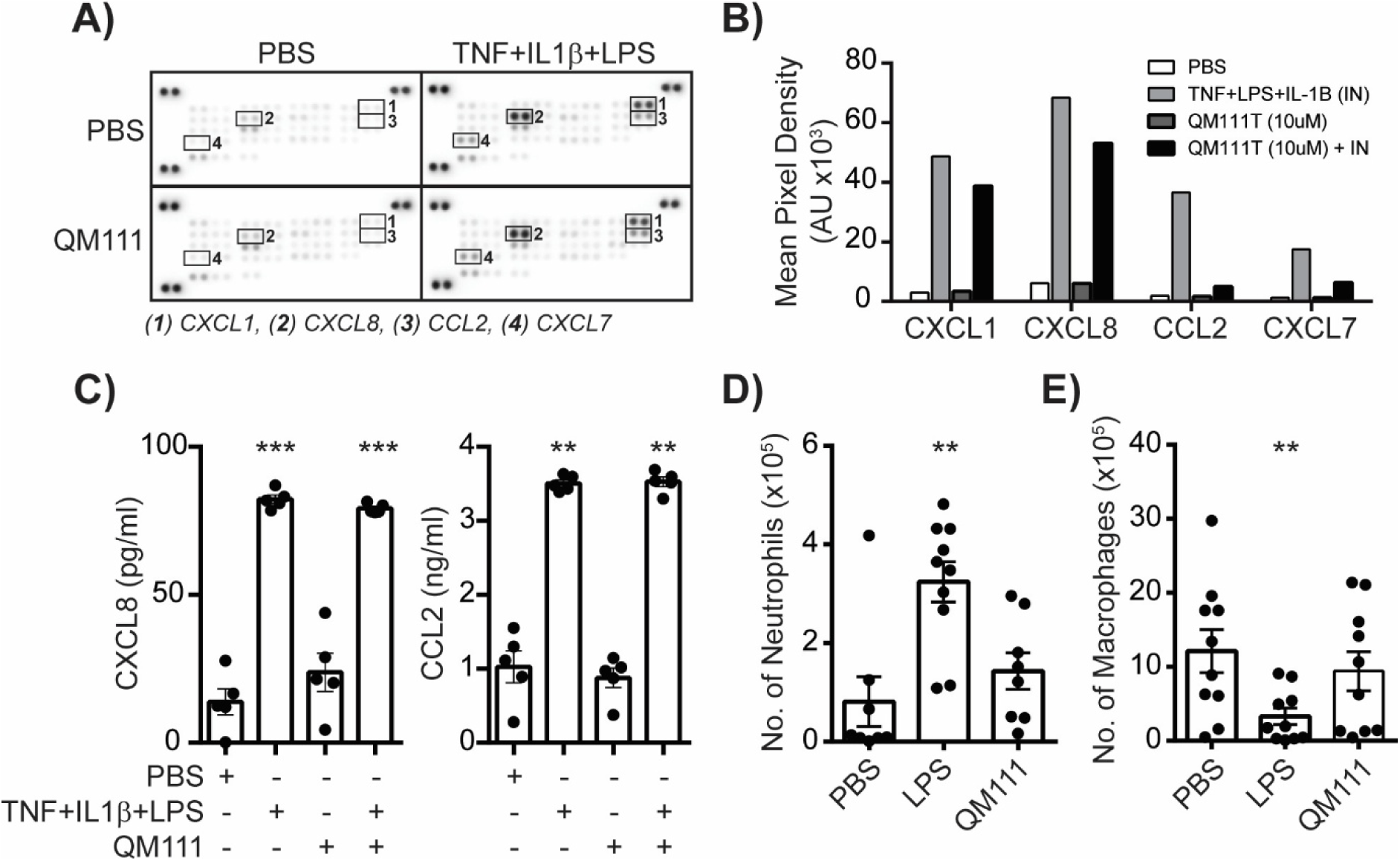
QM111 does not elicit an inflammatory response. **A)** Conditioned medium from HUVECs treated with either PBS, TNF + Il1β + LPS (inflammatory cocktail), QM111 (10µM) or QM111 + the inflammatory cocktail was assayed for a range of chemokines using the human proteome profiler chemokine array. **B)** Densitometry of the most prominent spots (CXCL1, CXCL8, CCL2 and CXCL7). **C)** CXCL8 and CCL2 levels were measured in conditioned medium of HUVECs (**p≤0.01, ***p≤0.001, n=5). **D)** Neutrophil and **E)** macrophage counts in murine peritoneal lavage 4h after LPS stimulation or treatment with QM111 (Statistical comparisons are to PBS control, **p≤0.01, n=∼9).

We next examined whether antibodies could be generated against the QM111 peptide sequence and whether pre-existing human IgG reactivity to the peptide was detectable in human sera. Rabbits immunised with KLH-conjugated QM111 in Freund’s adjuvant generated a strong antibody response, confirming that the ELISA could detect QM111-specific antibodies under positive-control immunisation conditions (**Figure 5A**, **Supplemental Figure 4A**). We then screened human sera from nine donors by ELISA using serial serum dilutions and anti-human IgG detection. No detectable IgG reactivity against QM111 was observed in the human sera tested (**Figure 5B**). No signal was detected against a scrambled QM111 control peptide in either human serum samples or anti-QM111 rabbit serum, supporting sequence-specific recognition of QM111 by the polyclonal antibody (**Figure 5C and D**).

**Figure 5:**
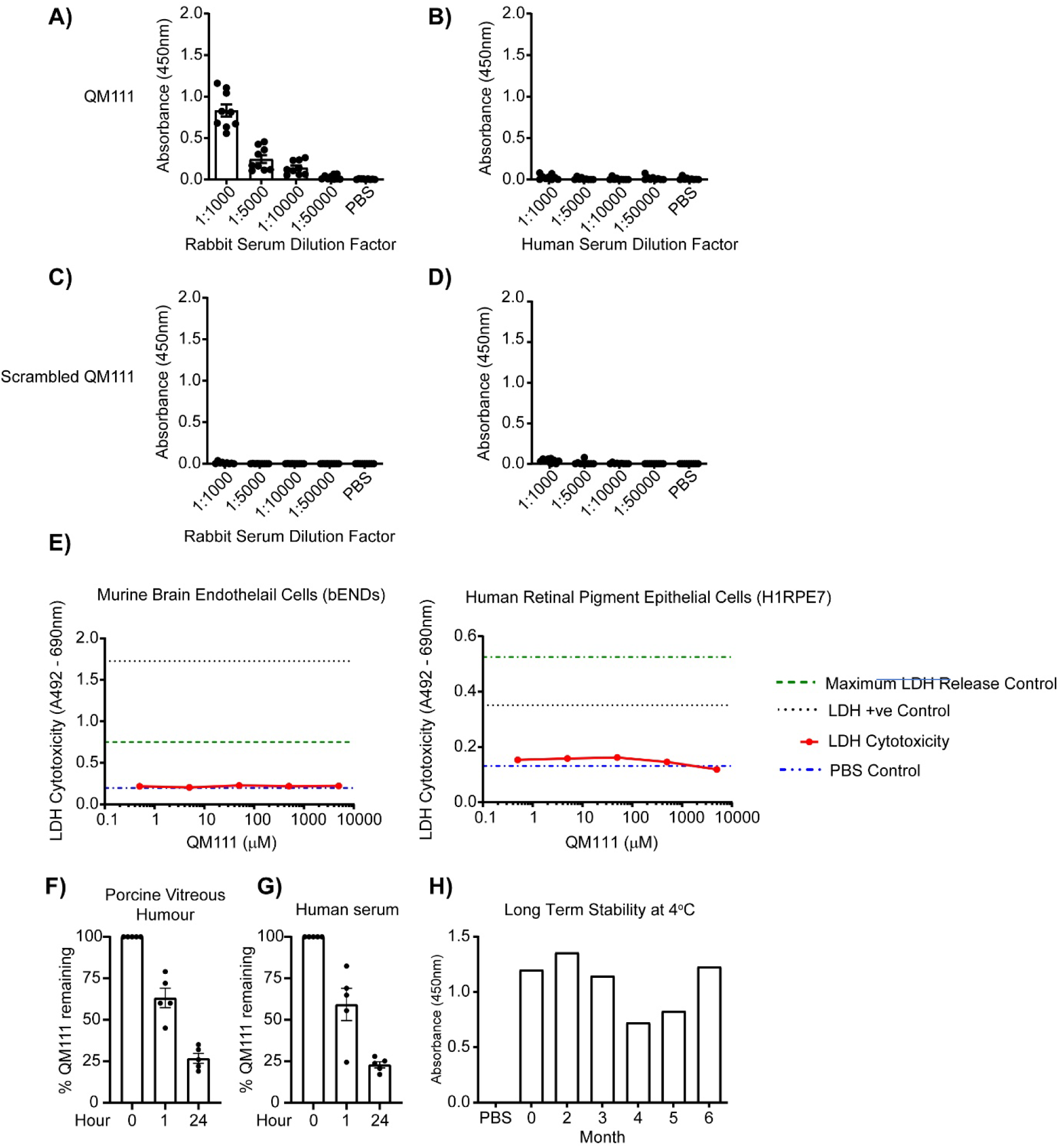
QM111 is not cytotoxic and has a good stability profile. **A)** Immuno-reactivity of rabbit serum after immunisation with KLH linked QM111 in the presence of Freund’s adjuvant. ELISA was performed with the indicated dilutions of QM111 antisera. **B)** Pre-existing human IgG reactivity to QM111 was not detectable. Serum from 9 human donors was assayed at the dilutions indicated. **C)** and **D)** neither QM111 rabbit antisera, nor human sera showed any immunoreactivity to a scrambled version of QM111. **E)** Measurement of lactate dehydrogenase was used as a measure of cytotoxicity in cells (either endothelial cells isolated from brain tissue derived from a mouse with endothelioma - bEND.3 cells - or human retinal pigment epithelial cells - H1RPE7 cells) treated with the indicated concentrations of QM111. **F)** QM111 (100 mM) was incubated in porcine vitreous humour at 37°C for the times indicated and the amount of peptide quantified by ELISA using the rabbit polyclonal antibody raised to the peptide sequence. **G)** Levels of QM111 were measured by ELISA in human serum after its incubation with human serum at 37°C for the times indicated (each point represents serum from a donor. **H)** Solutions of 1mM QM111 in PBS were stored at 4°C over a 6-month period, and then QM111 levels were measured using ELISA.

Finally, to assess cytotoxicity, LDH release assays were performed in murine brain endothelial cells and human retinal pigment epithelial cells treated with increasing concentrations of QM111. No cytotoxicity was observed in either cell type, even at the highest concentrations tested (**Figure 5E**).

To assess the translational potential of QM111, its stability was examined in biologically relevant environments. QM111 exhibited good stability in both porcine vitreous humour and human serum, with approximately 60% of peptide detected after 1 hour of incubation at 37°C (**Figure 5F and G**). Although further degradation occurred over 24 hours, approximately 20–30% of peptide remained detectable. In addition, QM111 retained stability during long-term storage at 4°C for at least six months (**Figure 5H**). These findings indicate that QM111 possesses favourable stability characteristics for an unmodified therapeutic peptide.

In a final set of experiments, we set out to compare the anti-angiogenic properties of the syndecan-2 derived anti-angiogenic therapeutic (QM107) with those of QM111. Both peptides inhibited cell migration and angiogenic sprout formation to the same extent (**Figure 6A and 6B**). Of relevance to a pathological situation sprout formation from murine choroid explants was also inhibited to the same extent with both peptides (∼50-60%, **Figure 6C**). Importantly, treatment with a combination of both peptides reduced the standard deviation and the variability of the response indicating the potential of developing a combination therapy comprising QM107 and QM111.

**Figure 6:**
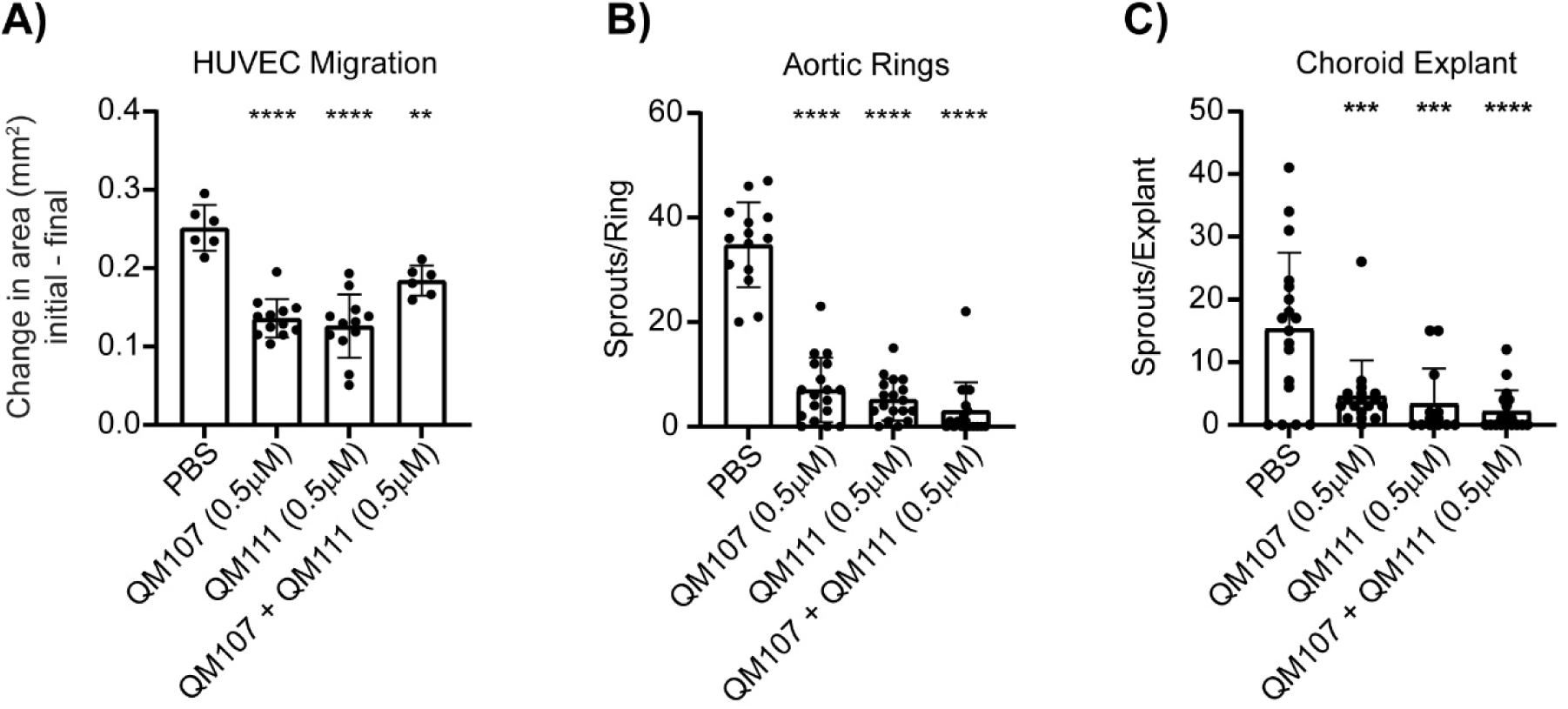
QM111 inhibits angiogenesis to the same degree as QM107 derived from SDC2. A) HUVEC migration in presence of QM111 and QM107, alone and in combination (n=7-14/condition, statistical comparison with PBS control, **p≤0.01, ****p≤0.0001). Sprout formation from aortic (B) and choroid (C) explants in presence of QM107, derived from syndecan-2, and QM111, derived from syndecan-3, and in presence of both (N=10-20, ***p≤0.001, ****p≤0.0001).

## Discussion

Syndecans represent an evolutionarily ancient family of cell surface proteoglycans. Invertebrates such as *C. elegans* and *Drosophila melanogaster* possess a single syndecan gene, as do primitive chordates including *Ciona savignyi* and *Branchiostoma lanceolatum*. Expansion to four family members occurred during vertebrate evolution and is conserved in mammals, birds and amphibians, whereas teleost fish have lost the syndecan-1 orthologue (Chakravarti & Adams, 2006). Syndecans from all these species all retain the same molecular architecture, an extracellular domain bearing glycosaminoglycan chains, a transmembrane domain and a short cytoplasmic domain (Gopal et al., 2016; Whiteford et al., 2008, Ricard-Blum and Couchman, 2023). The transmembrane and cytoplasmic domains of all these proteins are highly conserved whereas the extracellular domains are not, except for the sequences flanking the Ser-Gly GAG attachment motifs. Given the limited sequence conservation within syndecan ectodomains, it is remarkable that the four mammalian syndecans have been shown to regulate cell behaviour through discrete amino acid sequences embedded within their extracellular core proteins (Bertrand & Bollmann, 2019; De Rossi & Whiteford, 2013b). We report here the identification of a bioactive sequence of the SDC3 ectodomain that can inhibit angiogenesis and that a 9 amino acid sequence (P^195^-A^220^) of human SDC3 recapitulates the anti-angiogenic activities. We also provide a foundation for the development of novel peptide therapeutics for diseases characterized by pathological neovascularization. Taking together with previous studies of syndecan-1, −2 and −4, the present work establishes that all four mammalian syndecan family members contain discrete extracellular regulatory sequences capable of modulating cell behaviour.

The absence of sequence similarity between QM111 and previously identified syndecan-derived peptides (**Supplemental figure 5**), including synstatin and QM107 (Arokiasamy et al., 2025; Beauvais et al., 2009), suggests that each syndecan family member has evolved distinct regulatory modules that perform analogous biological functions through different molecular mechanisms. Despite the limited primary sequence conservation within syndecan ectodomains, functional conservation appears to have been maintained throughout evolution. We demonstrate that both the human and murine QM111 sequences retain comparable anti-angiogenic activity despite sequence divergence. Similarly, we have previously shown that the zebrafish syndecan-4 ectodomain supports cell adhesion in a manner comparable with its mammalian orthologue (Whiteford et al., 2008). Collectively, these observations suggest that evolutionary pressure has acted to preserve biological function rather than primary amino acid sequence within syndecan ectodomains.

Although the mechanism by which QM111 exerts its anti-angiogenic activity remains to be determined, its lack of sequence similarity to other syndecan-derived peptides suggests that it operates through a distinct molecular mechanism. Future studies should determine whether QM111 utilizes components of signalling pathways previously associated with other syndecan-derived regulatory peptides or instead defines a distinct endothelial signalling mechanism. Although regulatory peptides derived from syndecans 1, 2 and 4 frequently intersect with integrin-dependent signalling (Arokiasamy et al., 2025; Beauvais et al., 2009; Rossi et al., 2014; Whiteford et al., 2011), our preliminary attempts to define an integrin requirement for QM111 were inconclusive. This leaves open the possibility that QM111 acts through an integrin-associated pathway but also raises the possibility that the SDC3 sequence engages a distinct endothelial mechanism such as VEGFR2- or FGFR-mediated pathways.

Importantly, QM111 exhibited anti-angiogenic activity comparable with that previously described for QM107 issued from syndecan-2. Furthermore, combining QM111 with QM107 gave a more robust inhibition of angiogenesis, supporting the hypothesis that distinct syndecan-derived regulatory sequences may influence complementary signalling pathways. Whether this reflects engagement of different receptors or modulation of separate components of the angiogenic response remains to be determined.

An intriguing observation is that QM111, is predicted to reside within a region of comparatively reduced intrinsic disorder like the regulatory sequences within syndecan-1, and −2. Although these analyses are predictive, the localization of functional sequences to more ordered regions of the ectodomains of syndecans 2 and 3 raises the possibility that they contain conserved islands, which could fold in certain conformations or upon binding to a partner, embedded within an otherwise intrinsically disordered sequence. Whether this represents a general organizing principle underlying syndecan ectodomain function remains an intriguing question for future investigation. However, disorder prediction is dependent on the molecular context, and the isolated QM111 and QM107 peptides are 100% disordered whereas QM111M and QM111H are disordered at their N- and C-termini. In contrast to QM111 that does not contain protein binding regions predicted to undergo a disorder-to-order transition, the C-terminus of QM107 is predicted to be able to undergo such a transition. These features of the isolated peptides should be considered in future studies investigating their potential therapeutic applications.

An additional unresolved question is whether syndecan ectodomain regulatory sequences function in cis within the full-length receptor or in trans following ectodomain shedding. Synstatin has been proposed to function primarily in *cis* by disrupting molecular interactions at the cell surface, whereas the syndecan-2-derived peptide QM107 appears capable of acting in *trans* as a soluble effector following ectodomain shedding (Beauvais et al., 2009; Rossi et al., 2014). Whether QM111 behaves similarly remains unknown and may prove to be an important determinant of its physiological and therapeutic activity.

The identification of QM111 also highlights the therapeutic potential of endogenous bioactive sequences embedded within syndecan ectodomains. Here we have characterized the anti-angiogenic properties of QM111 in several different model systems. Importantly QM111 inhibits angiogenic sprout formation from murine choroid explants which is of direct relevance to neovascular eye disease. To determine whether QM111 could be a potential therapeutic, we performed comprehensive *in vitro* safety testing and observed negligible cytotoxicity, minimal inflammatory activity, an absence of preexisting antibody recognition in humans and favourable stability and storage characteristics. Given that QM111 is derived from an endogenous human protein, the minimal cytotoxicity, inflammatory activity and pre-existing antibody recognition observed in this study were perhaps unsurprising. Additionally, unlike many biologics, such peptides can be readily synthesized, modified and manufactured while retaining the potential to engage highly specific endogenous signalling pathways.

Collectively, these findings establish QM111 as a novel anti-angiogenic regulatory sequence within the SDC3 ectodomain and further support the concept that syndecan ectodomains represent evolutionarily conserved reservoirs of bioactive regulatory modules with significant therapeutic potential.

## Acknowledgements

JRW and GDR gratefully acknowledge funding from Arthritis Research UK (Grant No. 19207 and 21177), Fight for Sight (Grants 1558/59 and SGAFFS2203), Barts Charity (Grant No. MGU0313 and G-002397), Queen Mary Innovations, William Harvey Research Foundation, The Macular Society and the Dunhill Medical Trust (Grant No. RPGF1906\173). GDR is funded by a Diabetes UK (23/0006514) and Moorfields Eye Charity (GR001526) RD Lawrence fellowship.

## Author Contributions

**Samantha Arokiasamy**: Investigation, Methodology, Data curation, Formal analysis, Writing, Review & editing.

**Giulia De Rossi**: Investigation, Methodology.

**Thomas C. Mosely**: Investigation, Writing – review & editing.

**Sylvie Ricard-Blum**: Formal analysis, Methodology, Visualization, Writing – review & editing.

**James R. Whiteford:** Conceptualization, Methodology, Formal analysis, Supervision, Funding acquisition, Project administration, Writing – original draft, Writing – review & editing.

## Abbreviations

ANOVA: analysis of variance
BSA: bovine serum albumin
CCL: C-C motif chemokine ligand
CS: chondroitin sulphate
CXCL: C-X-C motif chemokine ligand
DAPI: 4′,6-diamidino-2-phenylindole
DMEM: Dulbecco’s modified Eagle medium
EC: endothelial cell
ECM: extracellular matrix
ELISA: enzyme-linked immunosorbent assay
FACS: fluorescence-activated cell sorting
FBS: fetal bovine serum
FGF2: fibroblast growth factor 2
GAG: glycosaminoglycan
GST: glutathione S-transferase
HS: heparan sulphate
HUVEC: human umbilical vein endothelial cell
IPTG: isopropyl β-D-1-thiogalactopyranoside
KLH: keyhole limpet hemocyanin
LDH: lactate dehydrogenase
LPS: lipopolysaccharide
PBS: phosphate-buffered saline
PCR: polymerase chain reaction
S3ED: syndecan-3 ectodomain
SDC: syndecan
SEM: standard error of the mean
TNF: tumour necrosis factor
VEGFA: vascular endothelial growth factor A
VEGFR2: vascular endothelial growth factor receptor 2

