## Supplemental material for "Identification of a novel anti-angiogenic regulatory sequence within the syndecan-3 extracellular core protein"

### SUPPLEMNTAL MATERIAL

Samantha Arokiasamy[1], Giulia De Rossi[2], Thomas C. Mosely[1], Sylvie Ricard-Blum[3] and James R. Whiteford[1*].

Affiliations:
[1]William Harvey Research Institute, Barts and The London School of Medicine and Dentistry, Queen Mary University of London, Charterhouse Square, London EC1M 6BQ, UK.
[2]Institute of Ophthalmology, University College London,11-43 Bath Street, London EC1V 9EL, UK.
[3]Institut de Chimie et Biochimie Moléculaires et Supramoléculaires (ICBMS), Villeurbanne, France.

Corresponding author:

*JRW, , William Harvey Research Institute, Queen Mary University of London, Charterhouse Square, London EC1M 6BQ, UK.

**Table 1: Primers used to generate deletion mutants**

S3EDA$^{45}$-A$^{194}$

For tttaaaggatccgctcaacgctggcgcaatg

Rev ttattaaggcttcgcctcgggagtgctag

S3EDP$^{195}$-L$^{380}$

For ttattaggatcccccccctgccacggctac

Rev tatataagcttctacagtatgctttctgagggag

S3EDE$^{92}$-V$^{310}$

For ttaattggatcccttgagacggccatgcgg

Rev tatataaagcttctgactggcacctctggctca

S3EDE$^{151}$-A$^{221}$

For ttaattggatccgaagagtccagccagaaag

Rev tatataaagcttctaagctgtggtcaggggaag

S3EDP$^{195}$-A$^{221}$

For ttaattggatccccccctgccacggctac

Rev tatataaagcttctaagctgtggtcaggggaag

**Supplemental Figure 1**

Human SDC1

A) 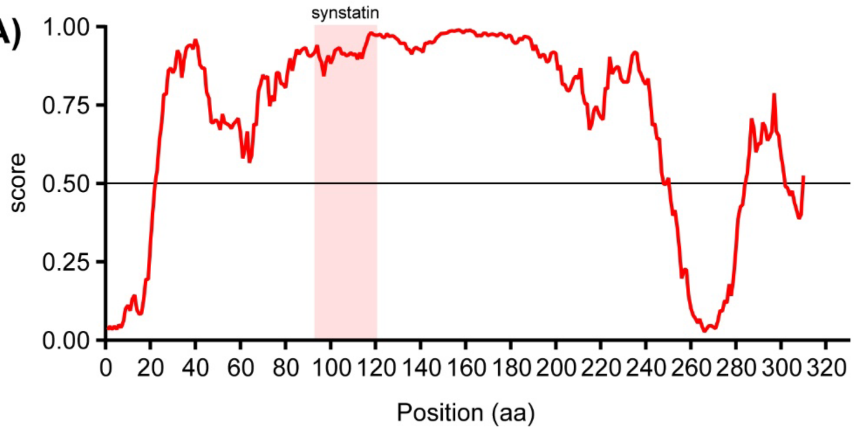


**Supplemental figure 1: A)** Predicted intrinsic disorder profile for the entire length of the human SDC1 amino acid sequence. The synstatin sequence is highlighted by a red box.

**Supplemental Figure 2**

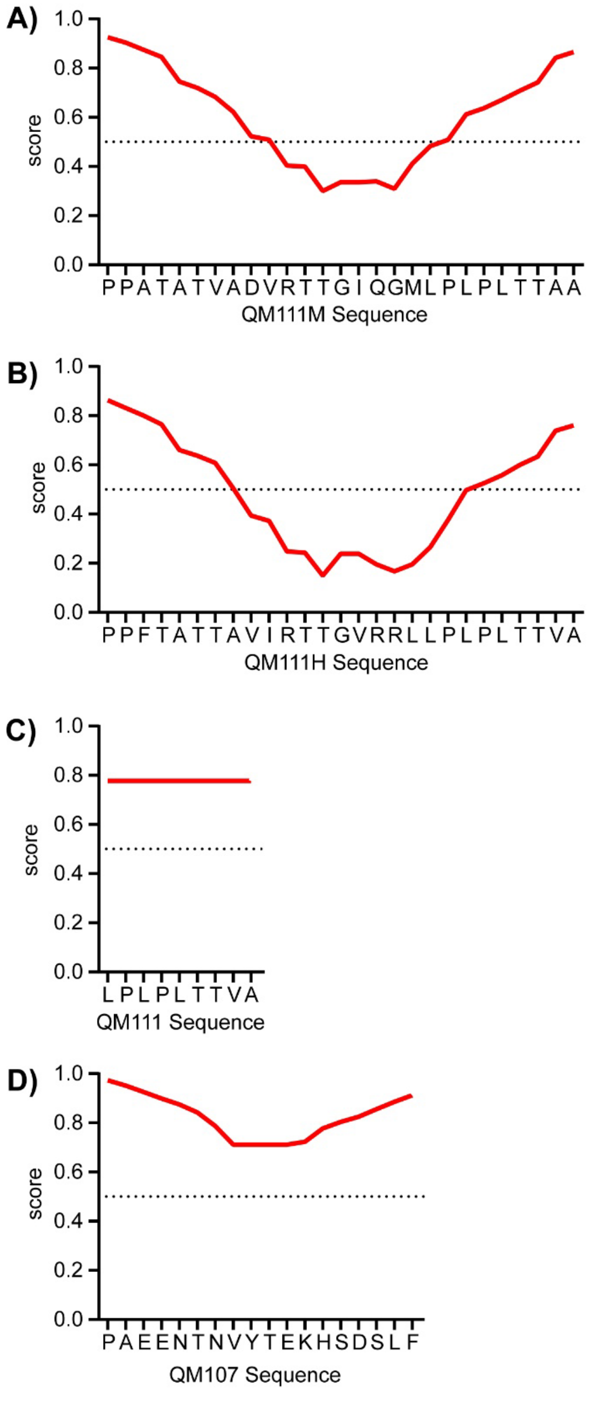


**Supplemental figure 2**: Prediction of intrinsic disorder in QM111 (A), QM111M (B) and QM111H (C) peptides of syndecan 3 and of QM107 (D) peptide of syndecan-2 using IUPred2A algorithm. Values above 0.5 indicate that the amino acid residue is predicted to be disordered. The numbering of amino acid residues starts with the first residue of the peptides.

**Supplemental Figure 3**

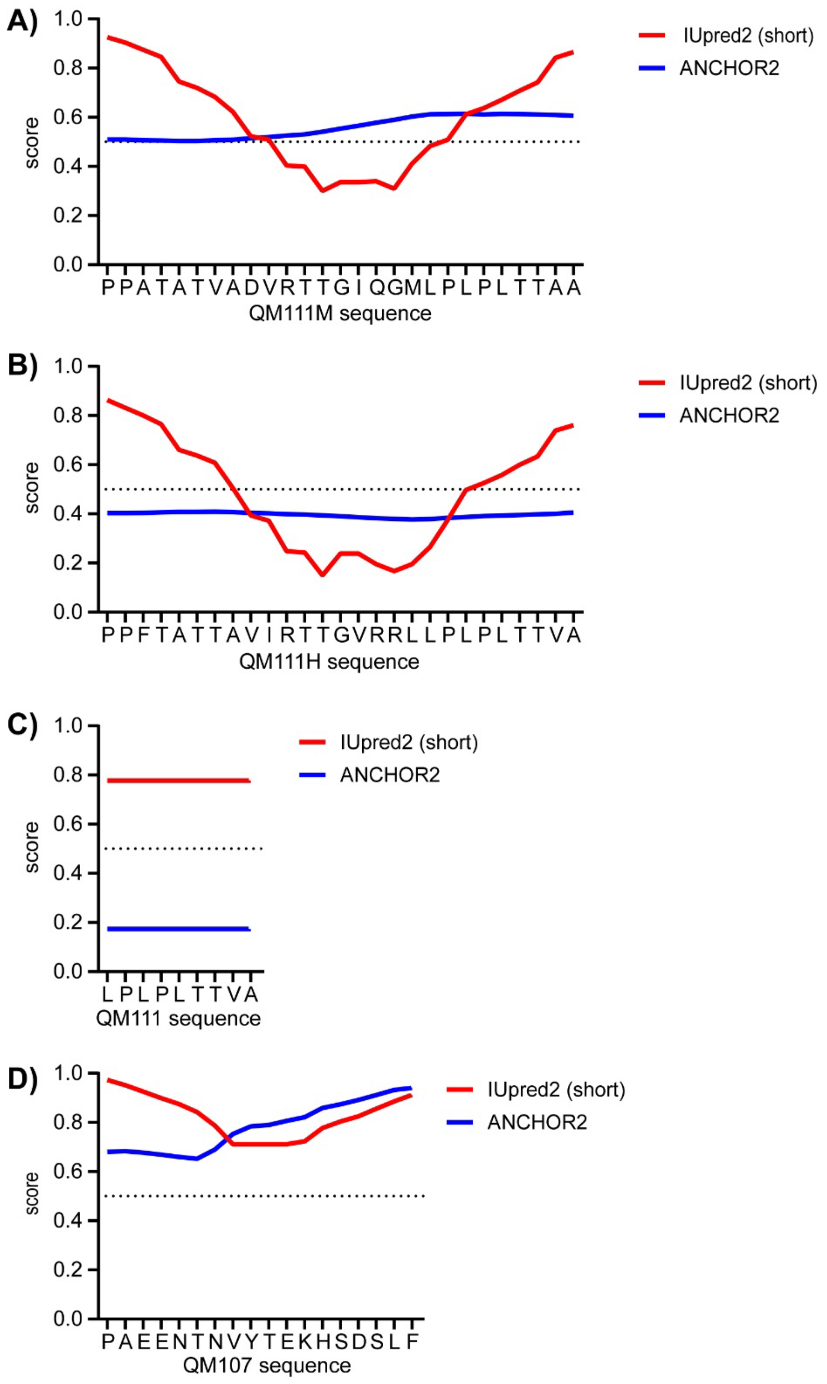


**Supplemental figure S3**: Prediction of intrinsic disorder using IUPred2A (red) and of disordered protein binding regions using ANCHOR2 (blue) in QM111 (A), QM111M (B) and QM111H (C) peptides of syndecan 3, and of QM107 (D) peptide of syndecan-2. Values above 0.5 indicate that the amino acid residue is predicted to be disordered

(IUPred2A) or that the sequence is predicted to be disordered protein binding region. The numbering of amino acid residues starts with the first residue of the peptides.

**Supplemental Figure 4**

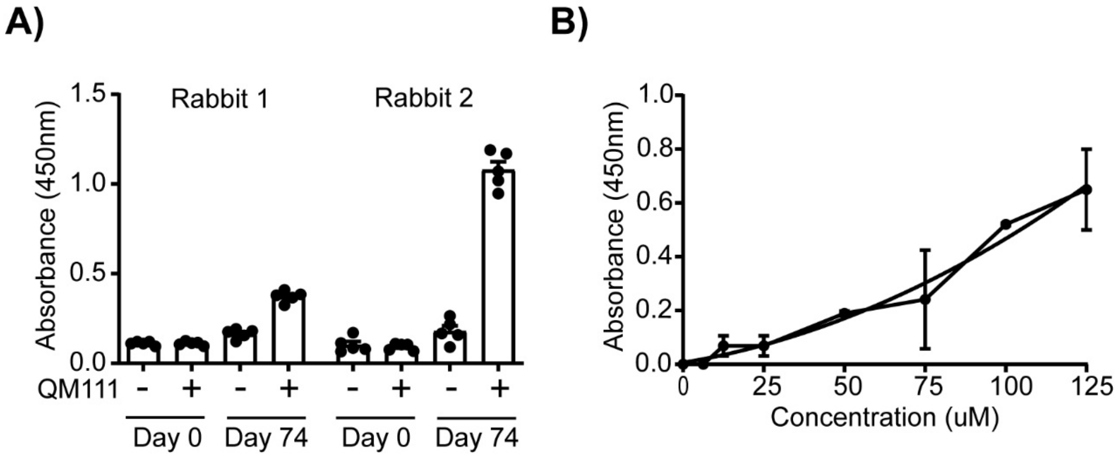


**Supplemental Figure 4: Generation of rabbit polyclonal antibodies to the QM11 peptide. A)** Comparison of immunoreactivity to QM111 of rabbit sera from the pre-immune bleed and 74 days post inoculation. Two rabbits were used and both showed strong immunoreactivity after 70 days whereas the pre-immunisation sera did not. **B)** Polyclonal anti QM111 sera show dose responsive immunoreactivity in the presence of human serum.

**Supplemental Figure 5**

**A)**

```
Synstatin(SDC1)         1 LPAGEGPKEAVVLPEVEPGLTAREQE 26
                          ||          ||....
QM111(SDC3)             1 LP----------LPTTVA--------  8

# Identity:       4/26 (15.4%)
# Similarity:     4/26 (15.4%)
# Gaps:          18/26 (69.2%)
# Score: -38
```

**B)**

```
QM107(SDC2)             1 PAEENTNVYTEKHSDSLF 18
                          .....|          :.
QM111(SDC3)             1 LPLPTT----------VA

# Identity:       1/18 ( 5.6%)
# Similarity:     2/18 (11.1%)
# Gaps:          10/18 (55.6%)
# Score: -34
```

**C)**

```
QM107(SDC2)             1 -PAEENTN--VYTEKHSDSLF----- 18
                           ||.|...  |...:....|.
Synstatin(SDC1)         1 LPAGEGPKEAVVLPEVEPGLTAREQE 26

# Identity:       5/26 (19.2%)
# Similarity:     6/26 (23.1%)
# Gaps:           8/26 (30.8%)
# Score: -33
```

**Supplemental Figure 5:** Sequence comparisons of syndecan bioactive peptides.
